# Sex differences in cell type-specific transcriptional programs and GPCR signaling in the human subgenual anterior cingulate cortex

**DOI:** 10.64898/2026.11.04.763991

**Authors:** Ariel W. Snowden, Niki Y. Li, Etienne Sibille, Max E. Joffe, Marianne L. Seney

**Affiliations:** Department of Psychiatry, University of Pittsburgh; Campbell Family Mental Health Research Institute, CAMH, Toronto; Departments of Pharmacology and Toxicology, and Psychiatry, University of Toronto; Department of Bioengineering, University of Pittsburgh

**Keywords:** subgenual anterior cingulate cortex, sex differences, inhibitory interneurons, transcriptomics, gene co-expression networks, electrophysiology

## Abstract

Psychiatric conditions show marked sex differences in disease onset, symptom presentation, and response to treatments. Understanding how sex differences manifest across molecularly distinct neuronal populations in individuals without psychiatric conditions may help explain sex-dependent variation in neural circuit function and treatment response. Here, we examined molecularly distinct cell types within the subgenual anterior cingulate cortex (sgACC) of non-psychiatric subjects. Using RNA sequencing data previously generated from laser-capture microdissected neurons, we investigated sex differences in gene expression and co-expression networks in excitatory pyramidal neurons (layers 2/3 and 5/6) and somatostatin (SST)-, parvalbumin (PVALB)-, and vasoactive intestinal peptide (VIP)-expressing inhibitory interneurons (N=10 males, 9 females). We identified sex differences in transcriptomic profiles, biological pathways, and co-expression network connectivity. PVALB cells showed stronger overall female transcriptional enrichment, while SST cells exhibited male-enriched transcriptional signatures. Weighted gene co-expression network analysis identified PVALB and SST modules more connected in males and females, respectively, which included hub genes involved in immune, extracellular matrix, and bioenergetic processes. Since G protein-coupled receptors (GPCRs) are viable therapeutic targets, we examined sex differences in GPCR transcripts. Across all GPCR-cell-type combinations, *CHRM2* in PVALB cells showed the second strongest sex difference (enriched in females). To test the functional relevance of this difference, we used patch-clamp electrophysiology in mouse prefrontal cortex. M2 receptor activation increased maximal firing rates in PVALB cells from female but not male mice. Collectively, our findings demonstrate cell type-specific sex differences in the sgACC corresponding to functional differences in neuromodulator signaling, with potential implications for treatment response.

## Background

The subgenual anterior cingulate cortex (sgACC) is a critical hub linking the prefrontal cortex with subcortical regions involved in emotional regulation^1,2^, reward processing^3,4^, and autonomic responses to emotional experiences^5^. The sgACC comprises areas 24 (a and b), 25 and 32, located ventral to the genu of the corpus callosum^6^. In primates, area 25 is connected to ventromedial and posterior orbitofrontal regions, temporal cortex, thalamus, amygdala, hippocampus, and ventral striatum^6–9^. These connections position the sgACC as an interface between higher-order cognitive control and subcortical emotion, arousal, and reward systems.

Glutamatergic pyramidal neurons and GABAergic interneurons govern information flow within these cortical circuits. Three major inhibitory interneuron populations can be defined by mutually exclusive expression of somatostatin (SST), parvalbumin (PVALB), or vasoactive intestinal peptide (VIP) cells^10^. These cell types generally subserve distinct microcircuit functions: PV cells gate pyramidal-cell spike-timing, SST cells regulate dendritic inputs, and VIP cells preferentially inhibit SST interneurons, thereby disinhibiting pyramidal cells. Inhibitory interneurons dynamically tune pyramidal cell excitation and help generate cortical oscillations that support higher-order cognitive functions, processes that are disrupted in psychiatric disorders^11–13^.

Alterations in these principal cell types in the sgACC have been examined in psychiatric disease^14–16^, but it remains unclear whether they exhibit sex-specific transcriptional profiles under baseline conditions. Psychiatric conditions show marked sex differences in incidence, onset, and treatment response. Among neurodevelopmental conditions, autism spectrum disorder is four times more prevalent in males^17^, while schizophrenia has an earlier age of onset in males than females^18,19^. In contrast, major depressive disorder has a twofold higher lifetime prevalence in women^20,21^. Determining whether there are sex differences in these cell populations in individuals unaffected by psychiatric disease may identify intrinsic molecular differences independent of disease and may help explain sex-biased disease vulnerability. Using a cell type-specific dataset of gene expression in the postmortem sgACC^16^, we examined sex differences in transcripts across inhibitory interneuron cell populations (PVALB, SST, VIP) and glutamatergic pyramidal cells. We identified distinct transcriptional profiles at the differential expression (DE), pathway, and network levels between males and females.

## Methods

### Human Subjects

Human brain samples were previously obtained from the University of Pittsburgh Brain Bank and the resulting RNA-sequencing data were published in Arbabi et al.^16^. The current study restricted all analyses to 19 nonpsychiatric control subjects (NPCs; female = 9, male = 10). Female and male subjects did not differ on age, pH, RNA integrity (RIN), postmortem interval, or intergenic mapping rate (see **S1 Table 1**). All procedures detailed in the original methods^16^ were approved by the University of Pittsburgh Committee for Oversight of Research and Clinical Training Involving Decedents and Institutional Review Board for Biomedical Research.

**Table 1.** Summary of sex-associated differentially expressed (DE) genes across sgACC cell types. For each cell type, the total number of differentially expressed genes (DEGs) and the number of male- and female-enriched DEGs are shown at two significance thresholds (*p* < 0.05 and *p* < 0.01), and an absolute log_2_FC > 0.26. Pink color indicates predominant female-enrichment; blue color indicates predominant male-enrichment.

| **Cell Type** | **Total DEGs (*p* < 0.05)** | **Male-enriched DEGs (*p* < 0.05)** | **Female-enriched DEGs (*p* < 0.05)** | **Total DEGs (*p* < 0.01)** | **Male-enriched DEGs (*p* < 0.01)** | **Female-enriched DEGs (*p* < 0.01)** |
|---|---|---|---|---|---|---|
| **PVALB** | 617 | 262 | 355 | 93 | 36 | 57 |
| **SST** | 693 | 461 | 232 | 148 | 115 | 33 |
| VIP | 571 | 350 | 221 | 96 | 66 | 30 |
| PyrL2/3 | 579 | 276 | 303 | 88 | 50 | 38 |
| PyrL5/6 | 591 | 286 | 305 | 105 | 59 | 46 |

### Cell Type Characterization and RNA-sequencing

Five cell types were identified for specific bulk transcriptomes: pyramidal layers 2/3 (PyrL2/3), pyramidal layers 5/6 (PyrL5/6), parvalbumin (PVALB), somatostatin (SST), and vasoactive intestinal peptide (VIP). The use of laser-capture microdissection (LCM) and RNAscope enabled anatomically and molecularly targeted collection of whole neuronal cell bodies from defined sgACC layers and interneuron populations, providing cell-type enriched transcriptomes containing both nuclear and cytoplasmic RNA. PVALB analyses included 18 subjects (9 per sex); insufficient gray matter precluded collection from one male subject. Detailed methods can be found in Arbabi et al.^16^ and in Supplementary Materials.

### Differential expression analysis

Following the preprocessing strategy detailed in Arbabi et al.^16^, gene filtering and normalization were performed separately for each cell type. Transcripts with >1 count in ≥ 80% of biological replicates were retained. Count data were normalized and transformed using voom^22^ to obtain log2 expression values and precision weights. These weighted expression measures were subsequently analyzed using linear models implemented using the limma^23^ package. Ensembl features were mapped to HGNC symbols before DE and WGCNA; features without an HGNC mapping were excluded, and only one Ensembl feature was retained per HGNC symbol. Separate linear models were constructed for each cell type (see **S1 Table 2**). The independent variable for all models was sex. Transcripts with *p* < 0.05 and log2 fold change > ± 0.26 were considered differentially expressed genes (DEGs). Log2 fold change values were calculated relative to females as the reference group, with negative values representing female-enriched genes and positive values representing male-enriched genes.

### Pathway representation

For each cell type, enrichment analysis was performed using Gene Ontology (GO) pathways in Metascape (https://metascape.org). Metascape groups related terms using hierarchical clustering, selects the most representative term per cluster, calculates enrichment scores, and visualizes pathway enrichment across cell types with heatmaps^24^. The resulting networks were imported into Cytoscape for visualization.

### Rank-rank hypergeometric overlap (RRHO)

To assess threshold-free overlap between the complete ranked DE results for each pair of cell types, we applied RRHO using the RRHO2 package^25^. RRHO computes a two-dimensional matrix of hypergeometric enrichment values across all possible rank thresholds, allowing for the identification of regions of significant concordant or discordant overlap without arbitrary DE cutoffs. For each pairwise comparison, gene lists were initially filtered to include only the genes present in both cell types. Within each list, genes were ranked on a continuous scale defined by RRHO score, calculated as -log_10_(p-value) multiplied by the direction of the effect size.

### Weighted Gene Correlation Network Analysis (WGCNA)

WGCNA was performed separately per cell type (WGCNA package in R^26^) to identify gene modules representing coordinated gene networks across donors. For each cell type, the top 5,000 most variable genes (based on variance across all NPC samples) were selected for module construction. Modules were created on the pooled sample (collapsed across sex) and preservation statistics were calculated to determine module robustness ^27^. Subsampling was done (50 iterations, 100 permutations), halving the sample to compare a test and reference networks to generate *Z* summary scores, where modules with a *Z* > 6, were selected for downstream module differential connectivity analysis.

#### Module Differential Connectivity (MDC) Analysis

MDC was conducted on the retained modules. MDC is the ratio of connectivity of all gene pairs within a module between males and females. An MDC >1 indicates greater connectivity in males than females, whereas an MDC < 1 indicates lower connectivity in males than females. We estimated the p-value based on 1,000 permutations of the following shuffling scheme: shuffled genes – adjacency matrix with random nodes but non-random connections. MDC values were considered statistically significant if *q* < 0.05, using the Benjamini-Hochberg procedure. Hub genes within a given network (and cell type) were identified using the ARACNe method^28^. If the n-hop neighborhood nodes (NHNN) for a given gene is greater than the mean NHNN + 1 SD, a gene is considered a hub. Sex-specific hubs were also identified (e.g., a gene that is only a hub gene in females for a given module). Cytoscape Version V3.10.4 was used to build and visualize hub gene networks.

### Mice

Mice were maintained on a 12:12 light-dark cycle (lights off at 7:00 a.m.) with food and water provided *ad libitum*. All mice were on a congenic C57BL/6J background and experiments were performed in adulthood (>10 weeks of age). Sex was limited to an assessment of external genitalia. Mice expressing tdTomato in PV-INs were generated by crossing female homozygous PV-Cre mice (Jackson Laboratories, Stock No. 017320)^29^ with male homozygous Rosa26-loxP-STOP-loxP-CAG-tdTomato mice (Jackson Laboratories, Stock No. 007909)^30^, yielding heterozygous PV-tdTomato offspring. All experiments were approved by the University of Pittsburgh IACUC and conducted in accordance with the NIH Guidelines for the Care and Use of Laboratory Animals.

### Brain Slice Preparation and Whole-Cell Patch-Clamp Electrophysiology

Electrophysiology experiments were performed as described^31,32^. Briefly, mice were anesthetized with isoflurane and decapitated. Brains were rapidly dissected and submerged in ice-cold N-methyl-D-glucamine (NMDG) cutting solution containing (in mM): 93 NMDG, 20 HEPES, 2.5 KCl, 0.5 CaCl_2_, 10 MgSO_4_, 1.4 NaH_2_PO_4_, 25 glucose, 5 Na-ascorbate, and 3 Na-pyruvate. Coronal slices (300 µm) through the prelimbic prefrontal cortex (PL-PFC) were prepared at room temperature in NMDG solution. Slices recovered for 10 min in warm (32–35°C) NMDG solution, then were transferred to room-temperature artificial cerebrospinal fluid (ACSF) containing (in mM): 119 NaCl, 2.5 KCl, 2 CaCl_2_, 1 MgCl_2_, 1 NaH_2_PO_4_, 11 glucose, and 26 NaHCO_3_ for at least 1 h prior to recording. All solutions were continuously oxygenated (95% O_2_/5% CO_2_).

Data were acquired using a Multiclamp 700B amplifier and pCLAMP 11 software (Molecular Devices). Slices were transferred to a recording chamber and perfused at 2 mL/min with heated ACSF (30–32°C). Borosilicate glass pipettes were pulled to 3–5 MΩ resistance (P-1000, Sutter Instruments) and filled with a potassium-based internal solution containing (in mM): 125 K-gluconate, 4 NaCl, 10 HEPES, 0.1 EGTA, 4 MgATP, 0.3 Na₂GTP, and 10 Tris-phosphocreatine. PV-INs in layer 5 of the PL-PFC were identified by tdTomato fluorescence, and cell identity was confirmed by the characteristic fast-spiking phenotype. After establishing whole-cell access, cells were dialyzed for 5 min under voltage clamp at −80 mV.

Membrane properties were assessed in current-clamp configuration. A series of 1-sec current injections were applied beginning at −150 pA, incrementing at 25 pA, and ending at +500 pA. All recording solutions contained the M1 mAChR antagonist VU0255035 (10 µM)^33^. Oxotremorine (OXO; 1 µM), an mAChR 1-5 agonist, was applied for 5 minutes, after which membrane properties were re-assessed. In some studies, the selective M2 muscarinic acetylcholine receptor antagonist AF-DX 116 (2 µM)^34^ was bath-applied for 5 min prior to and throughout OXO.

## Statistics

Data are presented as mean ± SEM. Statistical analyses were performed using GraphPad Prism. For electrophysiology experiments, *n* represents the number of cells and *N* represents the number of mice. For input-output curves, two-way mixed-effects analysis was used with Bonferroni post-hoc comparisons.

## Results

### Differential expression across cell types

We first compared gene expression between male and female NPC subjects in PyrL2/3, PyrL5/6, PVALB, SST, and VIP populations (**Fig.1**). In each neuronal subtype, multiple Y-linked genes, including *PCDH11Y*, *NLGN4Y*, *TTTY14*, and *USP9Y*, were enriched in males, while the X-linked gene *XIST*, which is required for X-chromosome inactivation, was strongly enriched in females. These sex-chromosome markers confirm that our analysis reliably captures meaningful sex differences in gene expression across all five sgACC cell types. Numerous female- and male-enriched DE genes were detected across cell types (**Table 1**). Among inhibitory interneurons, PVALB cells showed the strongest overall enrichment in females (n = 355 DEGs; **S2 Table 1**), whereas SST (n = 461 DEGs; Supplemental Table 2)and VIP cells (n = 350 DEGs; **S2 Table 3**) showed high enrichment in males. In excitatory pyramidal neurons, there were 303 female-enriched and 276 male-enriched genes in PyrL2/3 (**S2 Table 4**), and 305 female-enriched and 286 male-enriched genes in PyrL5/6 (**S2 Table 5**). These results indicate that sex differences in gene expression are detectable across all sgACC neuronal populations, but the magnitude and direction of these effects vary by cell type.

**Figure 1.**
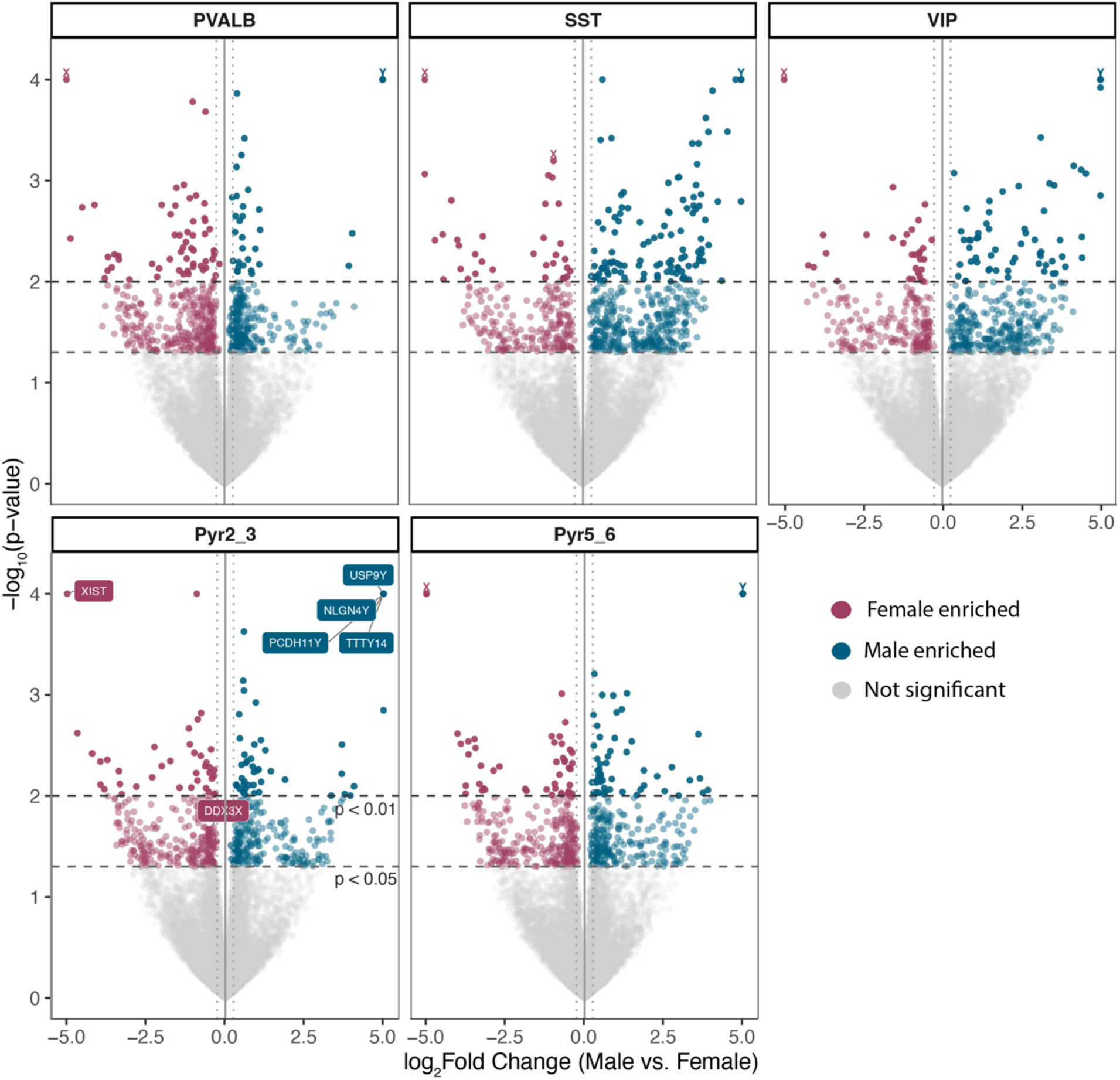
Volcano plots of differentially expressed genes between males and females in PVALB, SST, VIP, PyrL2/3, and PyrL5/6 cell types from nonpsychiatric control subjects. Log2FC plotted relative to -log_10_(*p*-value) by volcano plot. Horizontal dashed lines represent p-value significance cutoffs at p < 0.01 and p < 0.05. Blue circles represent Y chromosome genes and purple circles represent X chromosome genes.

### Sex differences in pathway enrichment across cell types

Pathway enrichment analysis revealed cell type-specific patterns of sex-biased biological processes (**Fig.2**). In PVALB cells, female-enriched transcripts mapped to synaptic and channel activity pathways, whereas male-enriched genes mapped to positive regulation of circadian rhythm, bone growth, and regulation of receptor-mediated endocytosis (**Fig.2A**). SST pathways included protein complex stabilizing activity and metabolic processes enriched in females and epithelial cell and tight junction processes enriched in males (**Fig.2B**). VIP cells showed only female-enriched pathways, including acetyltransferase activity, actin filament assembly, and phagocytic cup (**Fig.2C**). In PyrL2/3 pyramidal cells, female-enriched pathways included hydrogen peroxide metabolic process, catabolic processes, and asymmetric cell division, whereas male-enriched pathways included lymphoid progenitor cell differentiation and cation channel activity processes (**Fig.2D**). In PyrL5/6, female-enriched pathways included antigen receptor mediated signaling pathway and anaphase promoting complex binding, and male-enriched pathways included excitatory synapse, endothelial cell migration, and cell junction organization. Almost all pathways were uniquely enriched in either males or females in each cell type, except for the cell population proliferation pathway in PyrL2/3. These findings demonstrate limited convergence in sex-biased pathways across neuronal populations.

**Figure 2.**
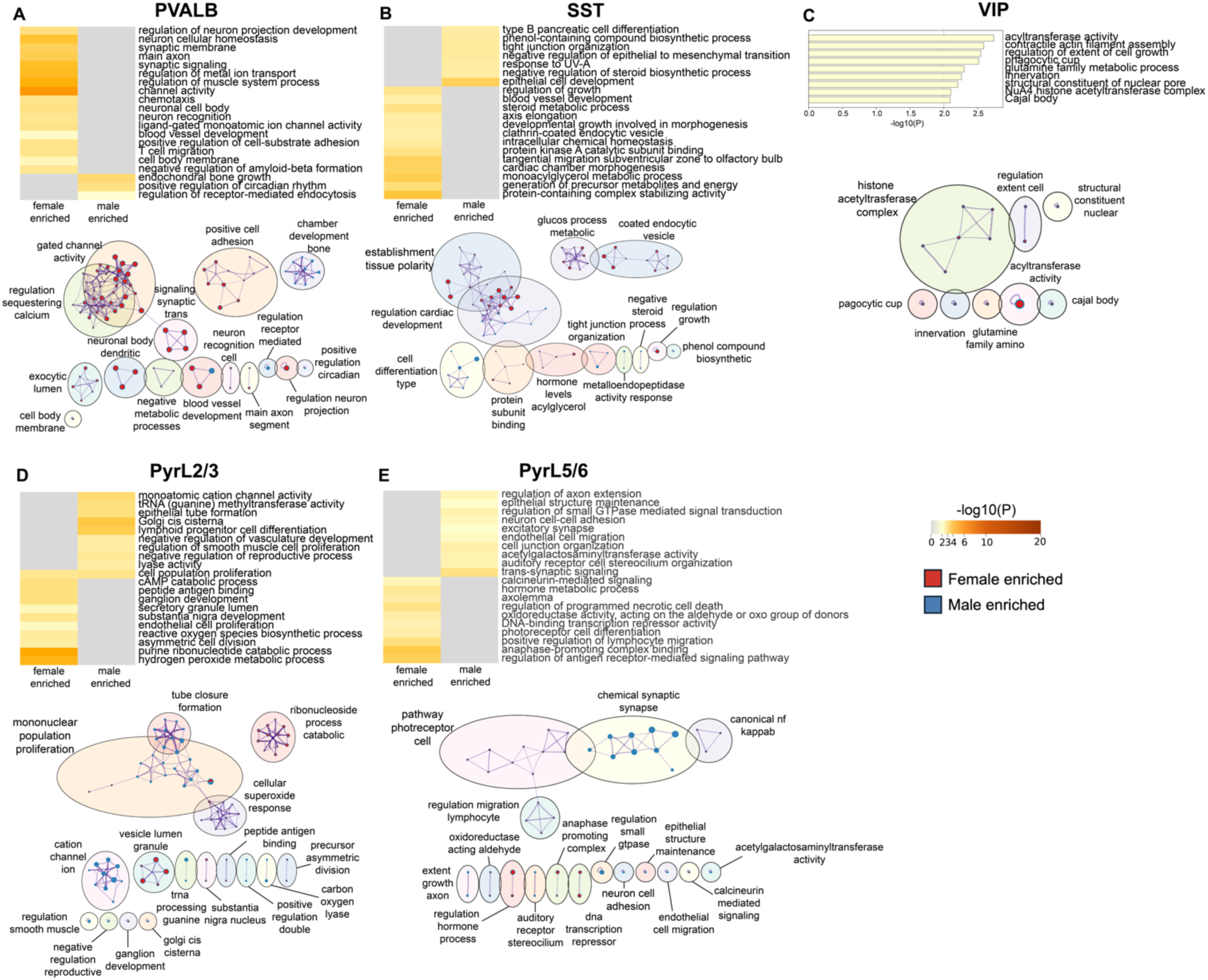
Sex differences in top enriched pathways across cell types. Heatmap and network visualizations of top biological pathways for female- and male-enriched genes in **(A)** PVALB, **(B)** SST, **(C)** VIP – pathways shown for VIP neurons were identified exclusively among female-enriched genes, **(D)** PyrL2/3, **(E)** PyrL5/6, using threshold *p* < 0.15 and log2FC > ± 0.26. Overlap of networks shows similarity among biological pathways. Nodes are sized proportionally to the number of associated genes and colored according to the direction of enrichment (female-enriched, red; male-enriched, blue).

### Pairwise RRHO comparisons reveal limited concordance across cell types

We next used threshold-free RRHO analysis of the complete ranked DE results to assess patterns across cell types. We asked whether any changes in gene expression were concordant (DE in the same direction together: e.g., up in both cell types or down in both cell types) or discordant (DE in opposing directions) across sgACC cell types. Cross-cell-type matrices exhibited little to no significant overlap between male-or female-enriched genes (**Fig.3**). This pattern suggests that the direction and magnitude of sex-associated DEGs are largely unique to each cell type. A modest signal was observed only between the pyramidal cell populations, which displayed hot spots in both the female-enriched (bottom left) and male-enriched (top right) quadrants. These findings collectively suggest that sex differences in transcriptional profiles are cell-type-specific rather than shared across cortical cell types.

**Figure 3.**
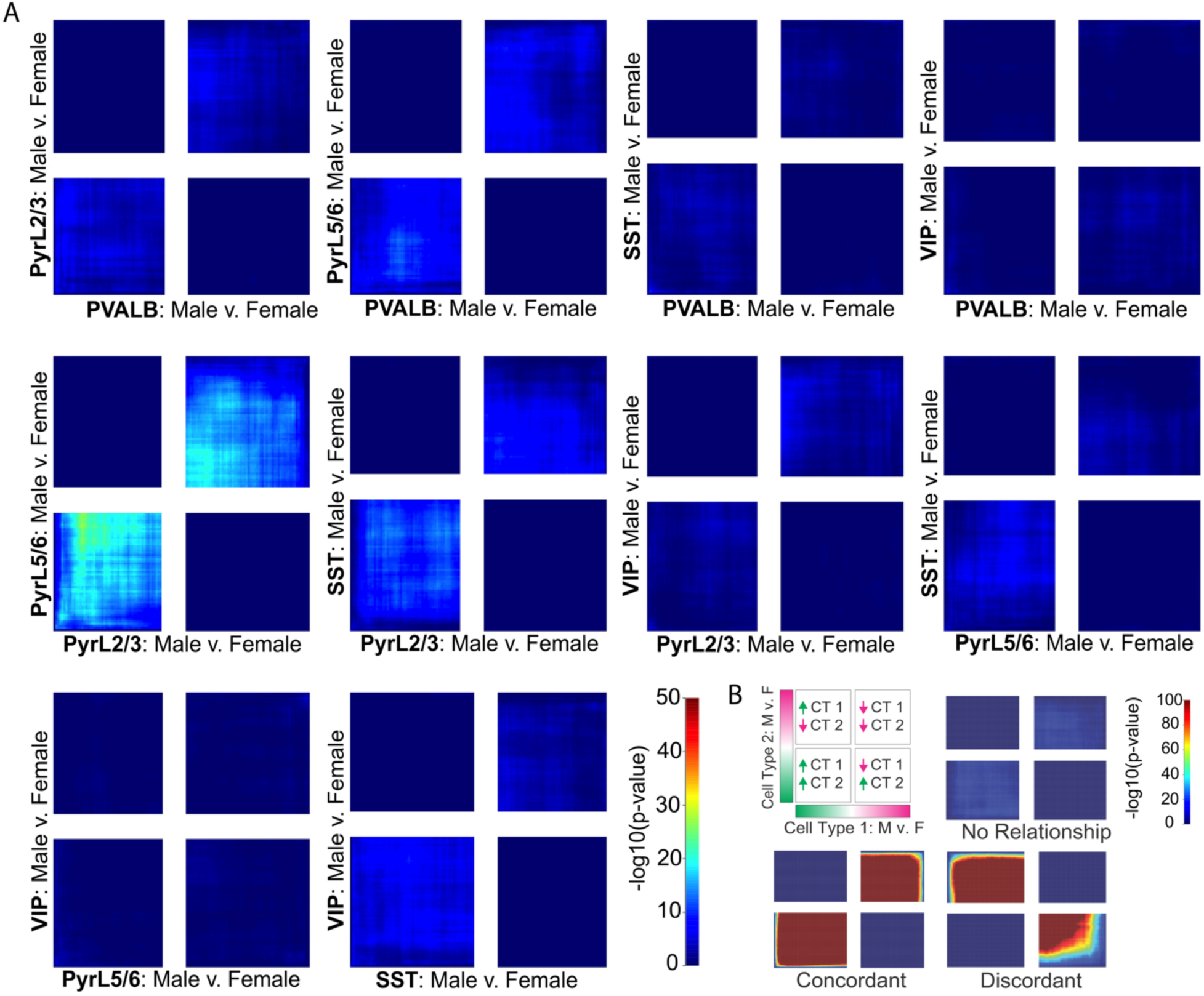
Rank-rank hypergeometric overlap (RRHO) comparisons between PVALB, VIP, SST, PyrL2/3, and PyrL5/6 cell types. **A)** RRHO analyses reveal minimal concordance or discordance across cell-types. Color scale reflects the -log_10_(*p*-value) with the maximum set to 50. **B)** Schematic illustrating RRHO interpretation, highlighting quadrants of concordant (shared direction of sex enrichment) and discordant (opposite direction of sex enrichment) gene overlap across cell types. CT = cell type.

### Sex-specific changes in connectivity of gene networks across cell types

For each cell type, we identified co-expression modules and asked whether the strength of connectivity within these modules varied by sex. We identified moderate to highly preserved modules (*Z_summary* > 6), with a total of 24 modules across all cell types that were retained for downstream MDC analysis to assess whether module connectivity changed based on sex (see **Fig.4A**). **Fig.4B,C** show clustering of genes in each module. An MDC ratio < 1, *q* < .05 indicates modules that are less connected in males relative to females, whereas an MDC ratio > 1, *q* < .05 indicates modules that are more connected in males v. females. For PVALB cells, 3 out of 4 modules were more connected in males than females (**S2 Table 6**). For SST cells, 3/4 modules were more connected in females than males (**S2 Table 7**). For VIP cells, 2/6 modules were more connected in females (**S2 Table 8**). For PyrL2/3 cells, 1 module was more connected in males (**S2 Table 9**); and for PyrL5/6 cells, 6/8 modules were more connected in males, and two in females (**S2 Table 10**). We focus on SST and PVALB modules because these cell types displayed the strongest male-specific and female-specific DE enrichment, respectively. Among the SST modules, the SST_1 module was more connected in females and exhibited the strongest enrichment for sex-associated differentially expressed genes (n = 145 genes, *MDC* = .095, *q* = .001) but yielded no significant pathway enrichment against the SST WGCNA gene universe. The SST_4 module was prioritized for hub network characterization because it combined significant DE enrichment and female-biased connectivity with broader functional coherence. Similarly, among PVALB cell modules, the PVALB_1 module was chosen due to having both significant connectivity differences and pathway enrichment. Therefore, these two modules of interest were selected for hub gene visualization: SST_4, which was more connected in females (n = 1517 genes, *MDC* = .661, *q* = .001) and significantly enriched for DE genes (*OR* = 1.67, *FDR-adjusted p* < .0001) and PVALB_1, a module with higher connectivity in males (n = 697 genes, *MDC* = 5.459, *q* < .001). Pathway enrichment indicated the top pathways in SST_4 were cytoskeletal motor activity and external side of plasma membrane (**Fig.4D**) and in PVALB_1 were response to ultraviolet light, signaling adaptor activity, T cell apoptotic process, and asymmetric synapse (**Fig.4E**).

**Figure 4.**
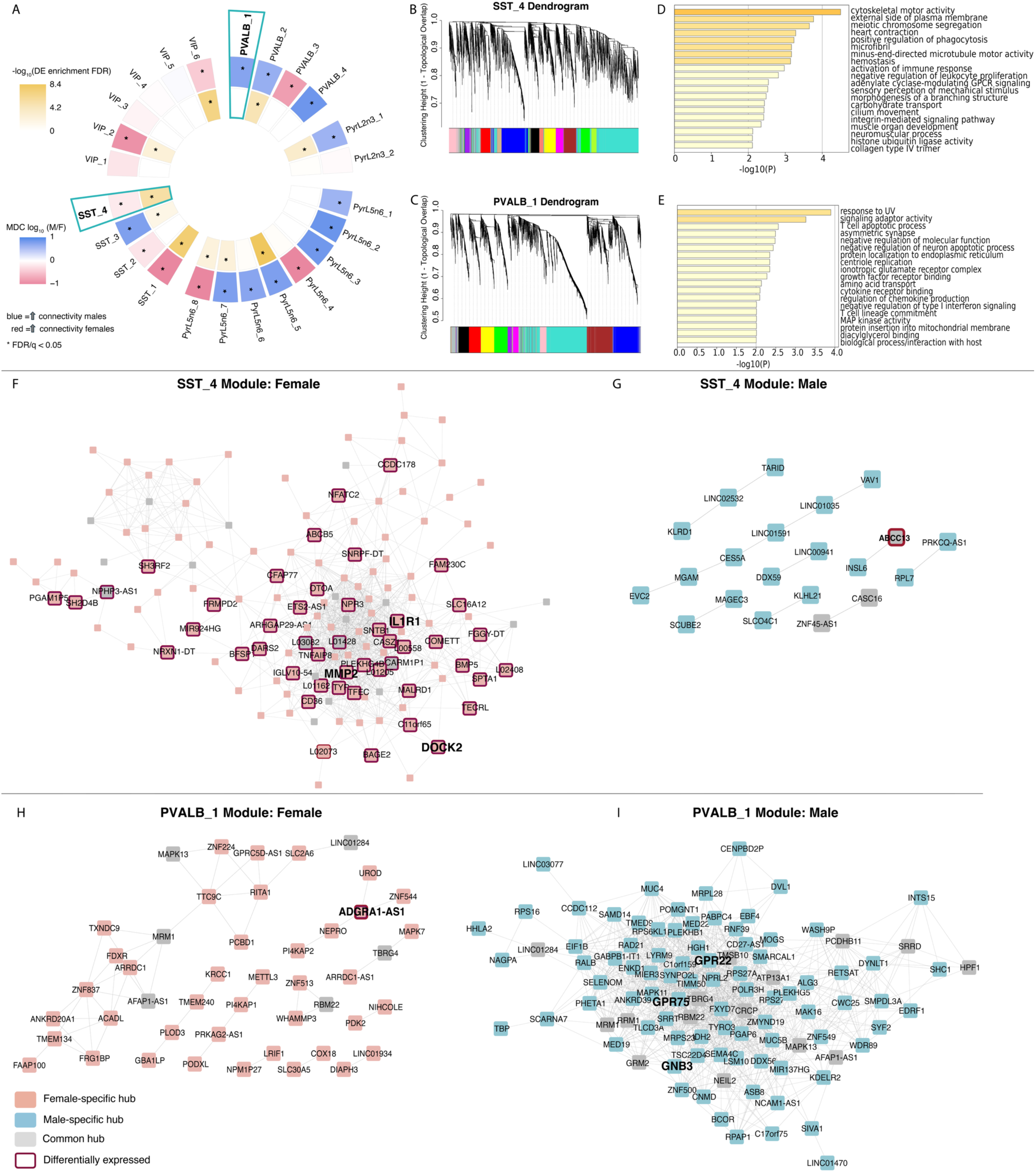
Weighted gene network correlation analysis reveals sex-specific co-expression changes across cell types. **(A)** Preserved WGCNA modules across five cell types (n=24). Inner and outer rings show DE-gene enrichment and sex-biased differential connectivity, respectively. Darker yellow indicates stronger DE enrichment; blue and pink indicate higher connectivity in males and females, respectively. SST_4 and PVALB_1 are outlined. **(B–C)** SST and PVALB module dendrograms. **(D–E)**. Module dendrograms for VIP, PyrL2/3, and PyrL5/6 are included in **S1** Figure 1. Pathway enrichment for SST_4 and PVALB_1, highlighting extracellular matrix-related processes. **(F–I)** Female and male hub-gene networks for SST_4 **(F–G)** and PVALB_2 **(H–I).** LINC genes in the SST_4 female hub network are abbreviated and begin with “L”. Nodes indicate female-specific (pink), male-specific (blue), or shared (gray) hubs; red outlines indicate DE genes. Edges represent co-expression correlations exceeding 97% for the SST_4 module and 90% for the PVALB_1 module.

### Targeted G protein-coupled receptor analysis highlights the strongest sex difference in expression of *CHRM2*

We aimed to test whether sex differences in gene expression translate into sex differences in protein function. We focused on GPCR expression, as this translationally relevant protein family represents the plurality of targets for medications, is readily accessible by small molecules, and modulates neuronal physiology in a rapid, cell-autonomous manner. With this goal in mind, we identified transcripts for GPCRs that exhibit sex differences in expression. Across all cell types, *GPR153* and *CHRM2* exhibited the strongest sex difference in gene expression, with higher expression in females compared to males (*GPR153* 3.2-4.2 logFC in SST and VIP cells; *CHRM2* 3.7 logFC in PV cells). No other GPCR exhibited >3 logFC. *CHRM2* codes for the M2 subtype muscarinic acetylcholine receptor (mAChR). There is well-developed pharmacology to enable functional experiments assessing the M2 receptor^35^, whereas GPR153 is an orphan receptor without any known ligands^36,37^. We therefore prioritized M2/*CHRM2* for subsequent functional studies.

### M2 receptor activation selectively increases PVALB cell maximum firing rates in female but not male mice

We obtained whole-cell patch-clamp electrophysiology recordings from fluorescently identified PVALB cells in prelimbic PFC of mice. We performed a series of injections to determine membrane properties of PVALB cells, then applied the muscarinic agonist oxotremorine-M (OXO; 1 µM) and repeated the current injection protocol (**Fig.5B-E)**. All recording solutions contained the M1 mAChR antagonist VU0255035 (10 µM)^33^. Recordings were made from slices of both female and male mice. An overall mixed-effects model revealed a sex difference in the effect of OXO (F_20,250_ = 1.6; p<0.05; current x sex x OXO interaction; **Fig.5F-G)**. Follow-up assessments revealed that OXO increased the maximum firing rate of PV cells in female mice, (F_3.4,25.5_ = 3.2; p<0.05; current x OXO interaction; **Fig.5F)**. By contrast, OXO did not affect spike-firing in PVALB cells from male mice (**Fig.5G)**. We next tested whether these actions were mediated by M2/*CHRM2*. Prior to OXO application, slices were incubated with the M2 antagonist AF-DX 116 (2 µM)^34^ (**Fig.5H)**. Under these conditions, OXO had no effect on current-evoked spiking in PVALB cells in female mice. Quantifying the change in maximum spike number for each cell confirmed that the OXO-evoked increase in PVALB interneuron firing in female mice was attenuated by AF-DX 116 pretreatment, (**Fig.5I**; F_3_ = 8.0, p<0.05, Kruskal-Wallis test; *:p<0.05 Dunn’s multiple comparisons test) confirming an M2-dependent mechanism. Together, these functional studies in mouse affirm functional relevance for the sex difference in *CHRM2* expression observed in PVALB cells in human sgACC.

**Figure 5.**
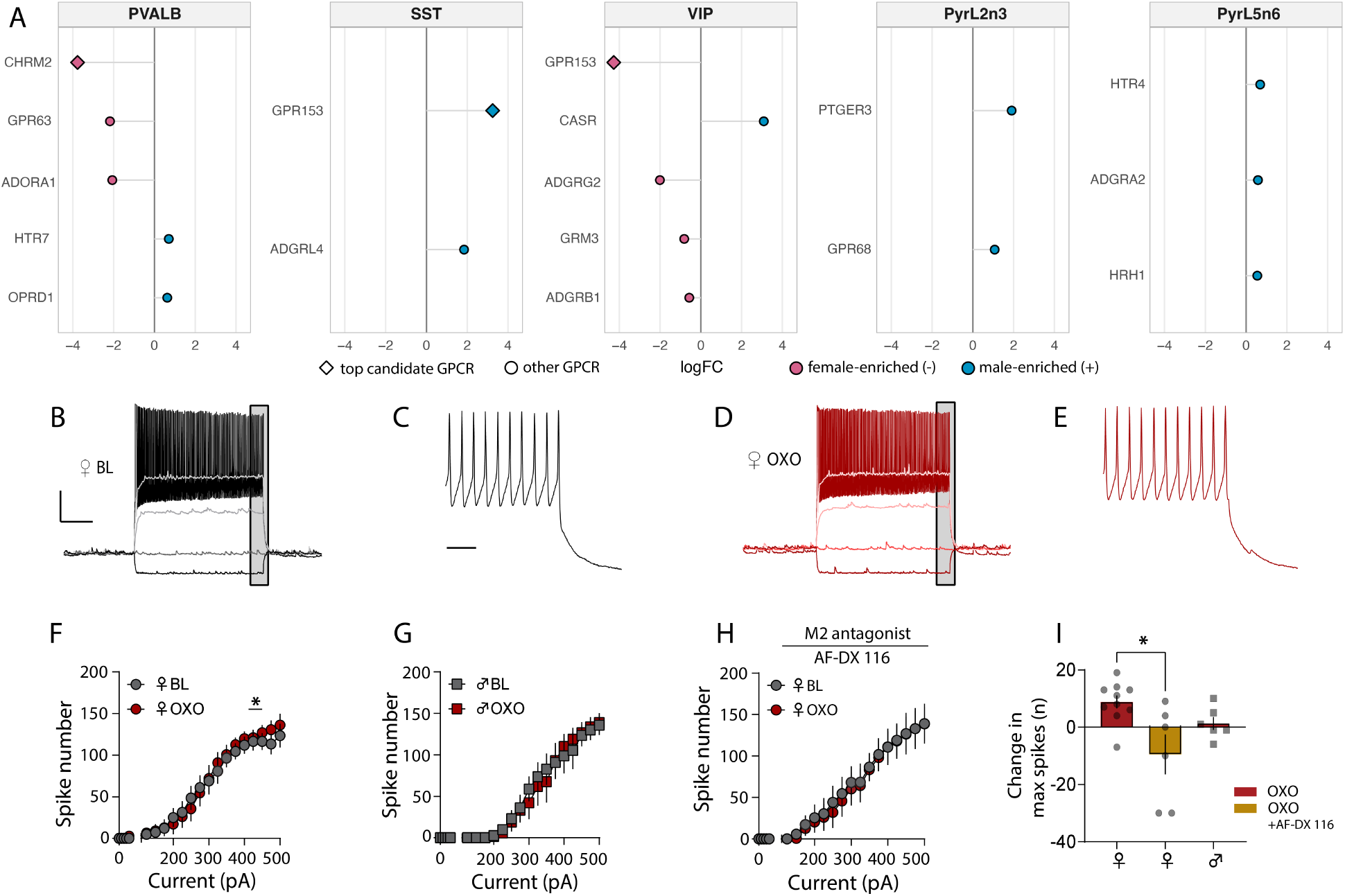
*Chrm2* is enriched in PVALB cells of the female human sgACC and drives sex-specific increases in PV cell firing via M2 receptor activation in the mouse PFC. **(A)** Top candidate GPCR genes across all five cell types. Negative log₂ fold change (pink-filled symbols) indicates enrichment in females, whereas positive log₂ fold change (blue-filled symbols) indicates enrichment in males. *Chrm2* showed one of the strongest log₂FC values, with enrichment in females relative to males. Whole-cell patch-clamp recordings from PFC PVALB interneurons. **(B-D)** Representative traces of recordings from a PV-IN from a female mouse. **(B)** Baseline (BL) responses to selected current injection steps (−100, 0, +100, +250, +400 pA). **(C)** Inset showing last 100 ms from +400 pA current injection. **(D)** Same as B, following incubation with the muscarinic agonist oxotremorine-M (OXO; 1 µM). **(E)** Same as C, following incubation with OXO. **(F-I)** Input-output curves depicting the maximum number of action potential spikes in response to a 1-second current injection (0–500 pA). All recordings were made in the presence of the M1 antagonist VU0255035 (10 µM). **(F)** In cells from female mice, OXO (dark circles) increased maximal spike-firing relative to BL (light circles) (F3.4, 25.5 = 3.3, p<0.05 current x OXO interaction; *: p<0.05 Sidak multiple comparisons test). **(G)** In male mice, OXO (filled dark red squares) did not alter spike-firing in response to current injections. **(H)** In the presence of the M2 mAChR AF-DX 116 (2 µM), OXO did not affect spike-firing in female mice. **(I)** Summarized findings showing the change in maximum spike number (OXO minus BL) across conditions. AF-DX 116 blocked the ability of OXO to increase maximum spike-firing in PVALB interneurons from female mice (p<0.02 Kruskal-Wallis test; *p < 0.05 Dunn’s multiple comparisons test). Individual data points represent single neurons. Data are presented as mean ± SEM. n/N = 6-10 cells / 3-5 mice per group.

## Discussion

Males and females showed distinct, cell-type-specific transcriptional profiles in the sgACC, with PVALB interneurons having more female than male-enriched DE genes, and these female-enriched genes mapped to synaptic and channel activity related processes. In contrast, SST interneurons had more male-enriched genes, which mapped to epithelial cell and tight junction related processes. Co-expression analyses further identified opposing changes in connectivity based in sex in PVALB and SST gene networks, highlighting an SST module more connected in females, and a PVALB module more connected in males. Hub genes identified in the SST network included immune signaling (*IL1R1, DOCK2*) and an extracellular matrix-related gene, *MMP2*. The PVALB network identified hub genes involved in GPCR signaling and synaptic processes. As GPCRs are compelling targets for treatment development, we followed up on one of the strongest female-enriched transcripts, *CHRM2* in PVALB cells. Oxotremorine, a non-selective muscarinic agonist increased PVALB cell excitability in female but not male mice, and we confirmed this effect was blocked by an M2 receptor antagonist. From the human transcriptome to rodent slice electrophysiology, our findings demonstrate that sex differences in the sgACC vary across inhibitory interneuron populations, and that sex differences in transcript expression can mediate functional effects on GPCR signaling.

Co-expression analysis identified sex-specific changes in network connectivity based in inhibitory interneurons in the sgACC. An SST-cell module (SST_4) that was more connected in females showed the strongest enrichment for DEGs. Pathway enrichment indicated that this module’s function was not a single canonical neuronal program, but rather a broader transcriptional state reflecting cytoskeletal motor activity, membrane-interface biology, and immune processes. Hub gene network analysis identified genes that may be key drivers of these processes. Although the module was more connected in females, many DEGs were enriched in males. Notably, we observed a 5-fold increase in *IL1R1* in males (logFC = 2.33). *IL1R1* (Interleukin 1 Receptor Type 1) encodes the primary cell-surface receptor for the pro-inflammatory cytokines IL-1α and IL-1β. Alterations in pro-inflammatory cytokines have been observed across multiple psychiatric conditions^38–40^. In schizophrenia, elevated *IL1R1* mRNA has been observed in the neurogenic subependymal zone, where it reliably predicts high-inflammatory profiles^41^. Intriguingly, *IL1R1* also regulates stress-responsive changes in gene expression in hippocampal SST and PVALB cells, including genes involved in synaptic organization, acetylcholine receptor signaling, and MAPK-related pathways^42^. This suggests a role for *IL1R1* in stress-induced synaptic alterations, raising the possibility that elevated *IL1R1* may prime male SST cells for altered inflammatory or stress-responsive signaling.

Other SST_4 hub genes pointed to similarly mixed sex-biased biology, including *DOCK2* and *MMP2*. The female-enriched *DOCK2* (Dedicator of Cytokinesis 2) has been investigated primarily in microglia and is involved in actin cytoskeletal remodeling required for immune-cell migration. In contrast, *MMP2* (matrix metallopeptidase 2), which encodes a zinc-dependent enzyme involved in cleaving extracellular matrix components, was male-enriched, but a female-specific hub gene. Collectively, these findings suggest that sex differences in SST cells may involve coordinated changes in inflammatory, extracellular matrix, and psychiatric disease-relevant signaling pathways.

Out of the four co-expression modules identified in PVALB cells, 3 modules were more connected in males, with 1 module more connected in females. For the module with the strongest connectivity in males relative to females, pathway enrichment indicated response to ultraviolet light, asymmetric synapse, and T cell apoptotic as top functional roles. Hub gene analysis revealed several components of GPCR signaling, including *GPR75*, *GPR22*, and *GNB3*. *GPR75* is co-localized to PVALB neurons in the mouse cortex^43^. The function of *GPR75* is best characterized in the context of metabolic regulation. Loss of function mutations in *GPR75* are linked to lower body weight and body mass index^44^ and *GPR75* knockout mice show resistance to weight gain during high fat diets^44^. Recent evidence attributes this metabolic function to *GPR75* expressed in glutamatergic neurons^45^. *GPR75* has also been shown to regulate mitochondrial function via AMPK signaling. In particular, higher *GPR75* expression in diabetic rodent models inhibit AMPK signaling, leading to mitochondrial dysregulation and the production of reactive oxygen species in hippocampal neurons^46^. GPR75’s potential role in sex-dependent metabolic dysregulation in psychiatric disease warrants further investigation. Additional hub genes support mitochondrial function including mitochondrial ribosome genes *MRPL28* and *MRPS23*, a mitochondrial protein import gene *TIMM50*, and *IDH2*, which contributes to mitochondrial metabolism. Together, these findings suggest convergence of GPCR signaling and energetic processes in a PVALB-cell module that was strongly connected in males but weakly coordinated in females. Further studies are needed to determine the functions of these GPCR-related genes in PVALB cells of the human sgACC.

In the human sgACC, *CHRM2* was the second strongest DEG among GPCR genes and selectively enriched in female PVALB cells. *CHRM2* encodes the M2 muscarinic acetylcholine receptor, a Gi-coupled receptor that canonically inhibits adenylyl cyclase and downstream cyclic AMP–protein kinase A signaling^47^. Using patch-clamp electrophysiology, we explored sex-specific effects of M2 receptor activation on PVALB cell firing in the mouse prelimbic PFC. In the presence of the M1 antagonist VU0255035, the muscarinic agonist oxotremorine-M increased maximum PVALB-cell firing in female but not male mice. This effect was abolished by the M2 antagonist AF-DX 116, supporting an M2-mediated effect on PVALB-cell firing in females. Prior work indicates that cortical PVALB interneurons can be more excitable in females than males and that this difference is regulated by GPCR signaling^32,48–51^. Manipulating PVALB-cell excitability also produces sex-specific effects on motivated and affective behaviors^50,52^. Our findings extend this work by identifying sex-specific effects of cholinergic signaling on PVALB-cell firing. Although increased firing was unexpected given the canonical inhibitory signaling of M2 receptors, other Gi-coupled receptors can enhance interneuron excitability. For example, activation of either D2 or D4 dopamine receptors increases the excitability of PVALB or fast-spiking interneurons in the rodent motor and medial prefrontal cortices^53–55^.

Muscarinic receptors have emerged as therapeutic targets for schizophrenia^56–58^ and major depressive disorder^59,60^, with evidence of sex differences in treatment response. The nonselective muscarinic antagonist scopolamine produces stronger antidepressant responses in women than men^59,60^. Although M1 receptors are generally emphasized, preclinical findings also implicate M2 receptors in antidepressant-like effects and associated BDNF/mTORC1 signaling^61,62^. A recent primate single-nucleus RNA-sequencing study further showed that ACC PV interneurons exhibit particularly high M2 expression and strong co-expression of M2 and M3 receptors^63^. Together, these findings support a role for sex-specific M2 signaling in the cholinergic modulation of cortical excitatory–inhibitory regulation and potentially treatment response in major depressive disorder.

The limited number of subjects of each sex reduced statistical power after correction for multiple testing; therefore, raw *p*-values were used for interpretation, as few findings survived FDR correction. Replication in larger samples using *p*-values corrected for multiple testing is needed.

Because our cell populations were identified using LCM sequencing, we were unable to detect distinct subpopulations within cell types. However, one advantage of LCM-seq is that it captures anatomically distinct whole-cell transcripts rather than only the transcripts present inside of the nucleus. Future studies should build on these strengths in larger, sex-balanced cohorts using LCM-seq to replicate the observed sex-related differences with adequate power for multiple-testing correction. Additionally, the use of a non-selective muscarinic agonist and an M2-preferring antagonist supports, but does not definitively establish, an M2-mediated mechanism. Future studies using genetic knockdown or deletion of *CHRM2* in PVALB cells would provide more conclusive convergent evidence.

## Conclusions

The current study analyzed transcripts from five cell types in the sgACC of male and female NPCs. We identified distinct transcriptional profiles based on sex and cell type, suggesting underlying sex differences in synaptic, immune, extracellular matrix, and bioenergetic processes in the absence of psychiatric disease. We further found that female-enriched *CHRM2* expression in human sgACC PVALB cells was accompanied by sex-specific changes in M2 function in rodents. Our preliminary findings represent a foundation for understanding sex-specific vulnerabilities to psychiatric disease.

## Supplementary Materials

**Supplemental Data 1 (S1):** Donor characteristics, differential expression model comparisons, and co-expression module dendrograms for additional cell types not shown in the main text.

**Supplemental Data 2 (S2):** Excel workbook containing cell-type-specific differential expression results (Tables 1–5) and module differential connectivity results and gene lists (Tables 6–10).

## Data Availability Statement

Newly generated electrophysiological data from the current study are available in the GIN repository: https:/doi.gin.g-node.org/10.12751/g-node.irk2dh/. Code for DE and WGCNA transcript analyses are available at https://github.com/devariel/sgACC-Cell-Type-Sex-Differences.git

## Notes

https://github.com/devariel/sgACC-Cell-Type-Sex-Differences.git

https:/doi.gin.g-node.org/10.12751/g-node.irk2dh/

### Competing Interest Statement

The authors have declared no competing interest.

### Summary of Updates

Updated the manuscript PDF to improve figure resolution and the clarity of figure labels and text.

### Funding

- National Institute on Drug Abuse: 1R01DA066210-01 (to MLS & MEJ)
- National Institute on Alcohol Abuse and Alcoholism: R00AA027806 (to MEJ)
- National Institute of Mental Health: 5T32MH016804-45 (training support for AWS)

