## Supplemental Data 1 for "Sex differences in cell type-specific transcriptional programs and GPCR signaling in the human subgenual anterior cingulate cortex"

**Supplementary Table 1. Demographic and technical variables by sex**

| **Variable** | **Female (n = 9)** | **Male (n=10)** | **Welch t (df)** | **p-value** |
| --- | --- | --- | --- | --- |
| Age, years | 48.33 ± 9.85 | 47.30 ± 11.47 | 0.21 (17.0) | 0.835 |
| Postmortem interval, hours | 17.37 ± 5.31 | 20.96 ± 4.85 | -1.53 (16.3) | 0.144 |
| RNA integrity number | 8.03 ± 0.70 | 8.03 ± 0.53 | 0.01 (14.8) | 0.991 |
| Brain pH | 6.67 ± 0.23 | 6.55 ± 0.22 | 1.14 (16.6) | 0.272 |
| Intergenic mapping rate | 0.22 ± 0.08 | 0.20 ± 0.04 | 0.43 (12.3) | 0.673 |
| Manner of death (Accidental) | 2 (22.2%) | 2 (20.0%) |  | 1.000 |
| Manner of death  (Natural) | 7 (77.8%) | 8 (80.0%) |  |  |
| Race (Black) | 1 (11.1%) | 0 (0.0%) |  | 0.474 |
| Race (White) | 8 (88.9%) | 10 (100.0%) |  |  |

**Note.** Continuous variables are summarized as mean ± SD. Categorical variables are shown as n (%).

Welch’s p value is used for continuous variables and Fisher exact P values are overall tests for each categorical

variable and are displayed on the first level only; blank cells do not indicate missing tests.

**Supplementary Methods**

*Cell Type Characterization and RNA-Sequencing*

Briefly, fluorescent *in situ* hybridization was performed on sgACC sections. Canonical markers were used to identify five cell types: pyramidal cells (*Slc17a7* marker), *Sst*, *Pvalb,* and *Vip*. A reference line located halfway between the plial surface and white matter was used to distinguish layers 2/3 from layers 5/6 pyramidal cells. Laser microdissection was used to capture 130 of each cell type from all subjects. RNA from all cell types was extracted using a PicoPure RNA isolation kit (Thermo Fisher Scientific, Waltham, MA) and libraries were prepared using SMARTer Stranded Total RNA-seq Pico Input Kit V2 (Clontech, Mountain View, CA). Quality control sequencing was performed using Illumina MiSeq sequencer (Illumina, San Diego, CA).

*Differential-Expression Model Comparison*

In Arbabi et al. (2025) Bayesian Information Criterion-based model selection identified sex as a predictor for all five cell types, along with cell-type-specific technical and demographic covariates. Because the present analysis was restricted to non-psychiatric control subjects, we evaluated whether intergenic mapping rate and manner of death remained necessary. For each cell type, the full model was compared with a reduced model excluding these covariates while retaining race where it had previously been selected. Gene-wise moderated F-tests were used to assess the contribution of the covariates removed from each model. Variance-partitioning analyses were performed using the **variancePartition** package (Hoffman & Schadt, 2016) to compare the proportion of expression variance attributable to sex under the full and reduced designs.

**Supplementary Table 2. Full and reduced differential-expression models by cell type**

| **Cell type** | **Full model** | **Reduced model** |
| --- | --- | --- |
| PVALB | ~ Sex + IntergenicRate + Race | ~ Sex + Race |
| SST | ~ Sex + IntergenicRate | ~ Sex |
| VIP | ~ Sex + IntergenicRate | ~ Sex |
| PyrL2/3 | ~ Sex + MOD | ~ Sex |
| PyrL5/6 | ~ Sex + IntergenicRate + Race + MOD | ~ Sex + Race |

**Supplementary Table 3. Covariate contributions and sex-associated variance in full and reduced models**

| **Cell type** | **Genes tested** | **Genes affected by covariate removal (nominal p < .05)** | **Sex variance: full model** | **Sex variance: reduced model** | **Difference (full − reduced)** |
| --- | --- | --- | --- | --- | --- |
| PVALB | 12,578 | 3.6% | 6.4% | 6.0% | +0.4 pp |
| SST | 12,193 | 10.0% | 5.8% | 5.3% | +0.5 pp |
| VIP | 13,161 | 11.2% | 5.7% | 5.3% | +0.4 pp |
| PyrL2/3 | 13,989 | 3.3% | 5.5% | 5.5% | −0.04 pp |
| PyrL5/6 | 13,746 | 3.6% | 5.9% | 6.1% | −0.15 pp |

**Note.** Full models retained the cell-type-specific covariates selected in Arbabi et al. (2025). Reduced models excluded intergenic mapping rate and/or manner of death (MOD) while retaining race where previously selected. The F-test column reports the percentage of genes for which the covariates removed from the reduced model had a nominal gene-wise p < .05. Sex-variance values were estimated using the variancePartition package. Positive differences indicate greater variance attributable to sex under the full model. Minor discrepancies between displayed variance percentages and differences reflect rounding. pp, percentage points. Based on these comparisons and to remain consistent with the original modeling strategy, full models were retained for all reported differential-expression analyses.

**Supplementary Results**

*WGCNA Module Dendrograms*

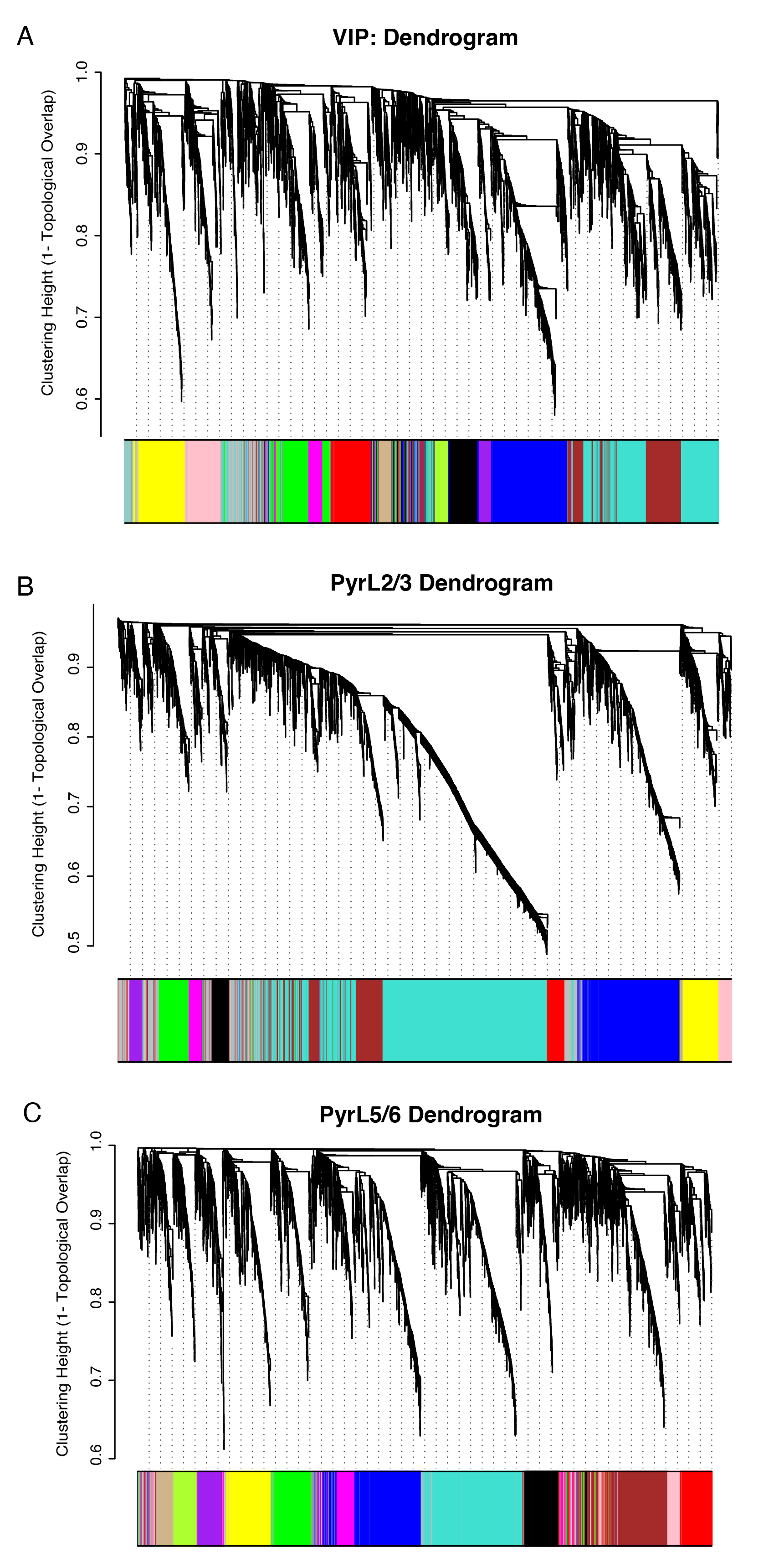

**Supplementary Figure 1. WGCNA Dendrograms.** Gene-clustering dendrograms and corresponding module assignments are shown for (A) VIP, (B) layer 2/3 pyramidal (PyrL2/3), and (C) layer 5/6 pyramidal (PyrL5/6) cells. Networks were constructed separately for each cell type using expression data pooled across male and female subjects. Each dendrogram branch represents a gene, and the color bar indicates its assigned co-expression module. Genes assigned to the same color exhibited similar co-expression patterns. Dendrogram height represents dissimilarity based on topological overlap.
